# Cavefish brain atlases reveal functional and anatomical convergence across independently evolved populations

**DOI:** 10.1101/832543

**Authors:** James Jaggard, Evan Lloyd, Anders Yuiska, Adam Patch, Yaouen Fily, Johanna E. Kowalko, Lior Appelbaum, Erik R. Duboue, Alex C. Keene

## Abstract

Environmental perturbation can drive the evolution of behavior and associated changes in brain structure and function. The generation of computationally-derived whole-brain atlases have provided insight into neural connectivity associated with behavior in many model systems. However, these approaches have not been used to study the evolution of brain structure in vertebrates. The Mexican tetra, *A. mexicanus*, comprises river-dwelling surface fish and multiple independently evolved populations of blind cavefish, providing a unique opportunity to identify neuroanatomical and functional differences associated with behavioral evolution. We employed intact brain imaging and image registration on 684 larval fish to generate neuroanatomical atlases of surface fish and three different cave populations. Analyses of brain regions and neural circuits associated with behavioral regulation identified convergence on hypothalamic expansion, as well as changes in transmitter systems including elevated numbers of catecholamine and hypocretin neurons in cavefish populations. To define evolutionarily-derived changes in brain function, we performed whole brain activity mapping associated with feeding and sleep. Feeding evoked neural activity in different sensory processing centers in surface and cavefish. We also identified multiple brain regions with sleep-associated activity across all four populations, including the rostral zone of the hypothalamus and tegmentum. Together, these atlases represent the first comparative brain-wide study of intraspecies variation in a vertebrate model, and provide a resource for studying the neural basis underlying behavioral evolution.

## Introduction

Brain function and behavior are influenced by evolutionary history and ecological environment [1, 2]. Robust differences in gross anatomy, neural connectivity, and gene expression have been associated with the evolution of behavaior in closely related species [3–5]. Most studies employing comparative anatomy have focused on a small number of brain regions, limiting insight into large-scale changes in brain structure and function. The recent generation of computationally-derived whole brain atlases and connectomes has provided an increased understanding of how neural circuits function [6–8]. These resources have largely been limited to a select number of genetically accessible model organisms, and have not been applied to a diverse set of models commonly used to study trait evolution. The generation of whole-brain atlases in closely related species, or even independent populations of the same species has potential to provide insight into the principles governing the evolution of brain structure and neural circuit connectivity associated with behavioral diversity.

The Mexican tetra, *Astyanax mexicanus*, comprises eyed surface fish that inhabit rivers throughout Mexico and at least 29 populations of cavefish in the San Luis Potosi region of northeast Mexico [9, 10]. Within the past 1 million years, multiple colonizations of caves by eyed surface ancestors has yielded independent cave populations that are geographically and hydrologically isolated from one another [11–14]. Cave populations of *A. mexicanus* have evolved numerous behavioral changes, including sleep loss, reduced social behaviors, wide-spread changes in sensory processing, and alterations in foraging behavior [17–21]. While trait evolution in cavefish has been studied for over a century, our understanding of brain evolution is largely limited to anatomical changes in a few brain regions including reduced size of the optic tectum and hypothalamic expansion in cavefish populations [15, 16]. Despite the long-standing focus on characterizing differences in behavior and morphology between cave populations, surprisingly little is known about the brain anatomy and neural circuits associated with these behavioral changes in cave populations.

Sleep and feeding are two homeostatically regulated behaviors that are essential for many aspects of biological function [22–25]. These behaviors interact at the genetic and neural circuit levels, and loss of sleep is associated with metabolism-related disorders [26, 27]. Although little is known about the genetic and evolutionary basis of these differences, it is hypothesized that the need for sleep is reduced in animals with greater foraging demands [28, 29]. We have previously identified the convergent evolution of sleep loss in multiple cavefish populations on sleep loss, cavefish displaying as much as an 80% reduction in sleep duration compared to surface fish counterparts [30]. In addition, feeding behavior differs dramatically between surface fish and cavefish. These differences include changes in the sensory modalities used to identify and capture prey in larval and adult fish, and hyperphagia in multiple adult cavefish populations [17,19,21]. Identifying the neural changes associated with the evolution of sleep and feeding behaviors may provide insight into fundamental principles governing the evolution of neural circuits and brain function.

Recently, the use of image registration to generate high-resolution reference brains has been used to map neural circuits in many model systems, from invertebrates through mammals [31–35]. In the zebrafish, multiple brain atlases have been developed that map neural circuits and putative connectivity between behaviorally relevant neurons [6,8,33,36]. These resources have provided unparalleled insight into brain function, but this approach has not been applied to study how evolution shapes brain development and function. Like zebrafish, larval *A. mexicanus* are transparent and amenable to whole-brain imaging in intact fish [37, 38]. In this study, we combined morphometric analysis, imaging of neural circuits, and whole-brain activity imaging to generate standard brains for river-dwelling surface fish and three independently evolved populations of cavefish. Using these reference brains, we quantified volume of defined brain regions and specific neuronal populations that contribute to sleep and feeding. We also mapped patterns of neuronal activity to the reference brain atlas to generate a brain-wide map of activity differences in waking, feeding, and sleeping fish using phospho-ERK labeling [6]. Together, these studies are the first comparative brain-wide analyses identifying differences in brain anatomy and function between populations with highly divergent behaviors.

## Results

### Behavioral and neuroanatomical evolution in larval *A. mexicanus*

To compare brain anatomy across independently evolved populations of *A. mexicanus*, we performed whole-brain confocal imaging in surface fish and Tinaja, Molino, and Pachón cavefish populations (Fig 1A). We first sought to determine whether differences in sleep and feeding phenotypes were present across all three cavefish populations at 6 days post-fertilization (dpf) when the fish are transparent, and their brains are accessible to intact whole-brain imaging. In agreement with previous findings at different developmental stages [39, 40], sleep is reduced across all three cave populations relative to surface fish (Fig 1B). The reduction in sleep is due to decreases in both total sleep bout number and average bout duration (and Fig S1). We previously found that the strike angle associated with prey-capture is increased in Pachón cavefish, likely due to an increased reliance on the lateral line in foraging behavior [17]. To test whether these evolved differences in prey capture are present in fish from other cave-dwelling populations, we measured strike angle in two other populations of cavefish. Strike angle was increased in Pachón and Tinaja, while Molino did not differ from relative to surface fish (Fig 1C).

**Figure 1.**
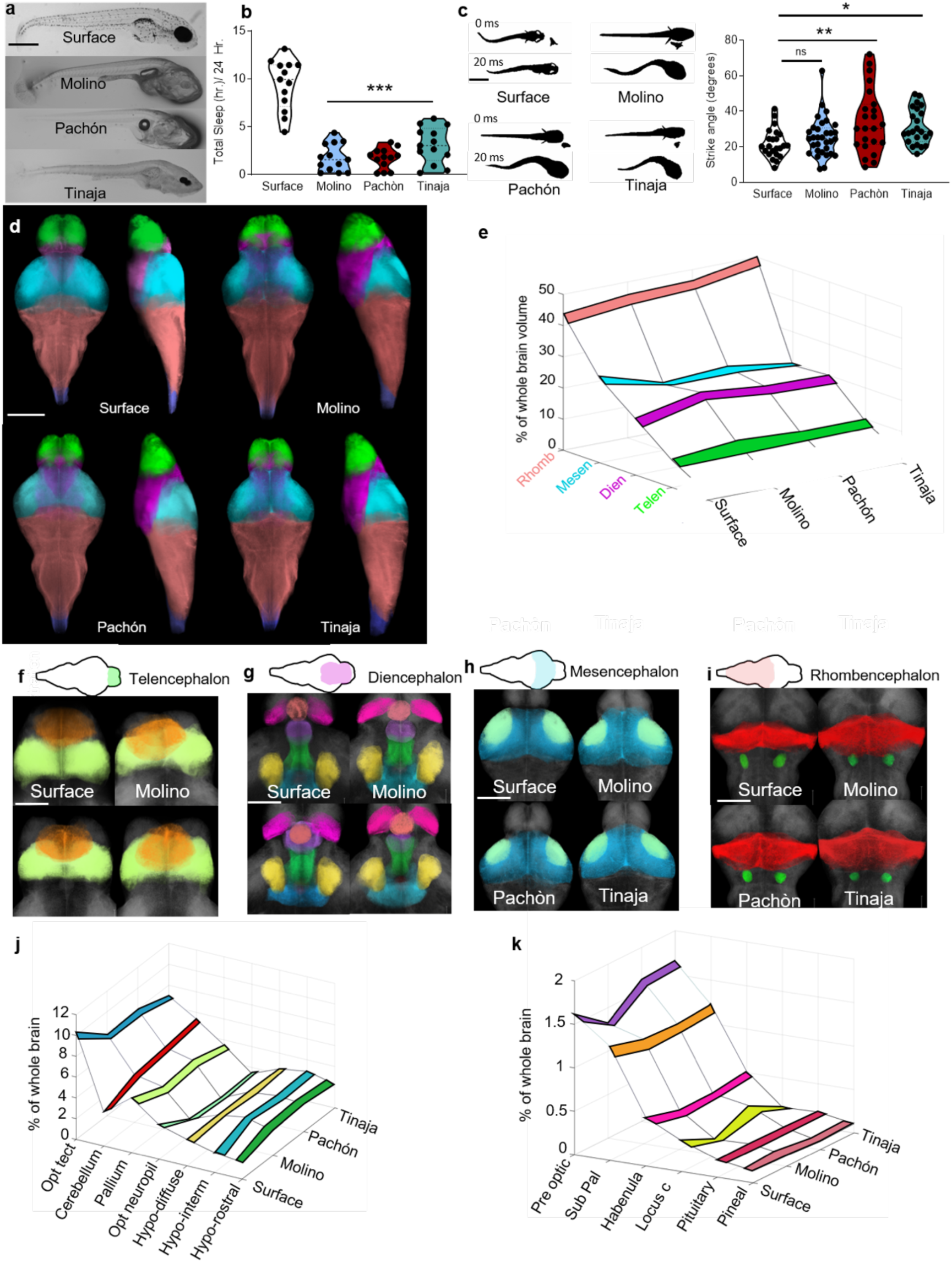
Behavioral and neuroanatomical evolution in larval *A. mexicanus*. **A.** Bright-field images of 6 dpf *A. mexicanus*: surface (top) and three cave-adapted populations: Molino (second from top), Pachón (second from bottom), and Tinaja (bottom). Scale bar, 500 µm. **B.** Total sleep over 24 hr in 6 dpf surface and cave populations (one-way ANOVA, F=51.53, P<0.0001, Dunnett’s multiple comparison test to Surface: Molino, p<0.0001 Pachòn, p<0.0001, Tinaja, p<0.0001). **C.** Feeding angle in larval *A. mexicanus*. Each population is shown while orienting to prey (0 ms) and then immediately after striking prey (20 ms) (one-way ANOVA, F=5.09, P=0.003, Dunnett’s multiple comparison to surface fish: Molino, p>0.56, Pachón, p<0.02, Tinaja, p<0.03). **D.** Anatomical segmentation of developmental regions in 6dpf brains using tERK antibody staining: telencephalon (green), diencephalon (magenta), mesencephalon (cyan), rhombencephalon (red), spine (blue). Scale bar = 300 µm. **E.** Quantifications of developmental regions segmentations normalized to whole brain size. All posthoc test were carried out comparing cavefish to surface fish. Telencephalon, (1-way ANOVA, F=0.845, P>0.47, Molino, p>0.46, Pachòn, p>0.35, Tinaja, p>0.68). Diencephalon, (1-way ANOVA, F=3.567, P<0.03, Molino, p<0.03, Pachòn, p<0.02, Tinaja, p<0.05). Mesencephalon, (1-way ANOVA, F=26.72, P<0.0001; Molino, p<0.001, Pachòn, p<0.001, Tinaja, p<0.001). Rhombencephalon, (1-way ANOVA, F=15.15, P<0.001; Molino, p<0.01, Pachòn, p<0.001, Tinaja, p<0.001) **F.** Volumetric projections of nuclei within telencephalon, including subpallium (orange), and pallium (light green) scale bar denotes 100 um. **G.** Volumetric projections of nuclei in diencephalon, including pineal gland (light red), habenula (pink), pre optic hypothalamus (purple), rostral zone of the hypothalamus (green), intermediate zone of the hypothalamus (blue), diffuse nucleus of the hypothalamus (yellow), and pituitary complex (dark blue). Scale bar denote 100 um **H.** Volumetric projections of nuclei within the mesencephalon. Optic tectum (blue), and optic neuropil (light green). Scale bar denotes 200 µm. **I.** Volumetric projections of nuclei within the rhombencephalon, showing cerebellum (red), and locus coeruleus (green). Scale bar denotes 200 µm. **J-K.** Quantifications of segmentations in F–I. For detailed information and statistics about all regions, see Table 1. N>12 for all sleep behavior, n>25 for all strike angle, and n>10 for all neuroanatomical segmentations.

Therefore, multiple independently evolved populations of cavefish have convergently evolved differences in sleep and feeding behaviors that manifest as early as at 6 dpf.

To characterize differences in brain anatomy between *A. mexicanus* populations, we quantified the size of individual brain regions within each *A. mexicanus* population. Immunostaining for total extracellular signal–regulated kinase (tERK) has been established in zebrafish as a method for labeling broad anatomical regions within the central nervous system [6]. At 6 dpf, we immunolabeled larvae for tERK and obtained whole-mount confocal images through the entire brain. tERK signal was detected throughout the brain in all *A. mexicanus* populations tested (Fig S2A), confirming that tERK immunostaining also serves as a broad neuroanatomical marker in *A. mexicanus*. Total brain volume did not differ between surface fish and fish from the three cave populations (Fig S2B). We therefore quantified the size of individual brain regions, normalized to the volume of the entire brain. Volumetric quantification revealed convergence on changes in the major brain subdivisions that are established during neurodevelopment across all three cave populations relative to surface fish. The rhombenchephalon and the diencephalon were expanded and the mesencephalon was reduced in fish from all three cave populations relative to surface fish (Fig 1D,E, Fig S3 and Table S1). These observations suggest that broad changes in brain structure are shared across independently-evolved cavefish populations.

To determine if there are changes in the size of brain regions that may be associated with evolved behavioral differences, we quantified 13 additional brain regions, including the tectum, cerebellum, pallium, and four regions of the hypothalamus (Fig 1F–K and Table S1, Fig S4, Movie S1, and Movie S2) in accordance with previously described nomenclature [7]. Consistent with previous reports, the optic tectum and neuropil were reduced, and the total hypothalamus volume was enlarged in all three cavefish populations [15,41,42]. The increase in hypothalamus volume was due to an enlargement of rostral and intermediate zones of the hypothalamus, with no differences between surface fish and cavefish populations in the volume of the diffuse nucleus of the hypothalamus (Fig 1K, Fig S4). The volume of the pineal gland, a region associated with secretion of sleep-promoting melatonin, was significantly reduced in all three populations of cavefish (Fig 1K, Fig S4), raising the possibility that these changes are associated with loss of sleep and circadian regulation of activity in cavefish [39, 43]. In addition, volume of the habenular nuclei that regulate stress were reduced in all three cavefish populations, demonstrating a potential neuroanatomical mechanism underlying blunted response to stress in cavefish (Fig 1K, Fig S4) [44]. While many of the evolved changes in brain anatomy we identified were shared between cave populations, we also observed differences between individuals from different cave populations. For example, the preoptic hypothalamus was reduced only in Molino cavefish, whereas the size of the locus coeruleus was significantly reduced in Molino and Pachón (Fig 1J and Fig S4). Together, these data suggest that cavefish from different populations have repeatedly evolved many of the same neuroanatomical changes in behaviorally-relevant brain regions.

### Neural circuitry associated with sleep and feeding

The circuitry underlying sleep/wake regulation is highly conserved across vertebrate species [45, 46]. Studies in zebrafish have identified a central wake-promoting role for the catecholamines dopamine and norepinephrine, and hypothalamic neurons expressing hypocretin/orexin (Hcrt) that consolidate wakefulness [47–50]. We previously reported that functional differences in β-adrenergic and HCRT signaling contribute to sleep loss in Pachón cavefish [18, 51], but the role of these signaling pathways in sleep regulation, and the neuroanatomy of catecholamine and HCRT neurons has not been characterized in other populations of cavefish. To determine whether specific classes of catecholamine neurons differ between *A. mexicanus* populations, we immunolabeled brains for *tyrosine* hydroxylase (TH), and quantified TH+ neurons throughout the brain (Fig 2A). The number of TH+ neurons in the locus coeruleus, a highly conserved wake-promoting region, did not differ between any of the *A. mexicanus* populations we examined (Fig 2B). TH+ neurons were more abundant in the telencephalon of both the Pachón and Tinaja populations, but the number of neurons in the telencephalon did not differ between Molino and surface fish (Fig 2C). Furthermore, the number of TH+ cells in the pretectal area of the brain was significantly reduced in all cavefish populations (Fig 2D), and hypothalamic TH+ neurons were more abundant in all three populations of cavefish than in surface fish (Fig 2E).

**Figure 2.**
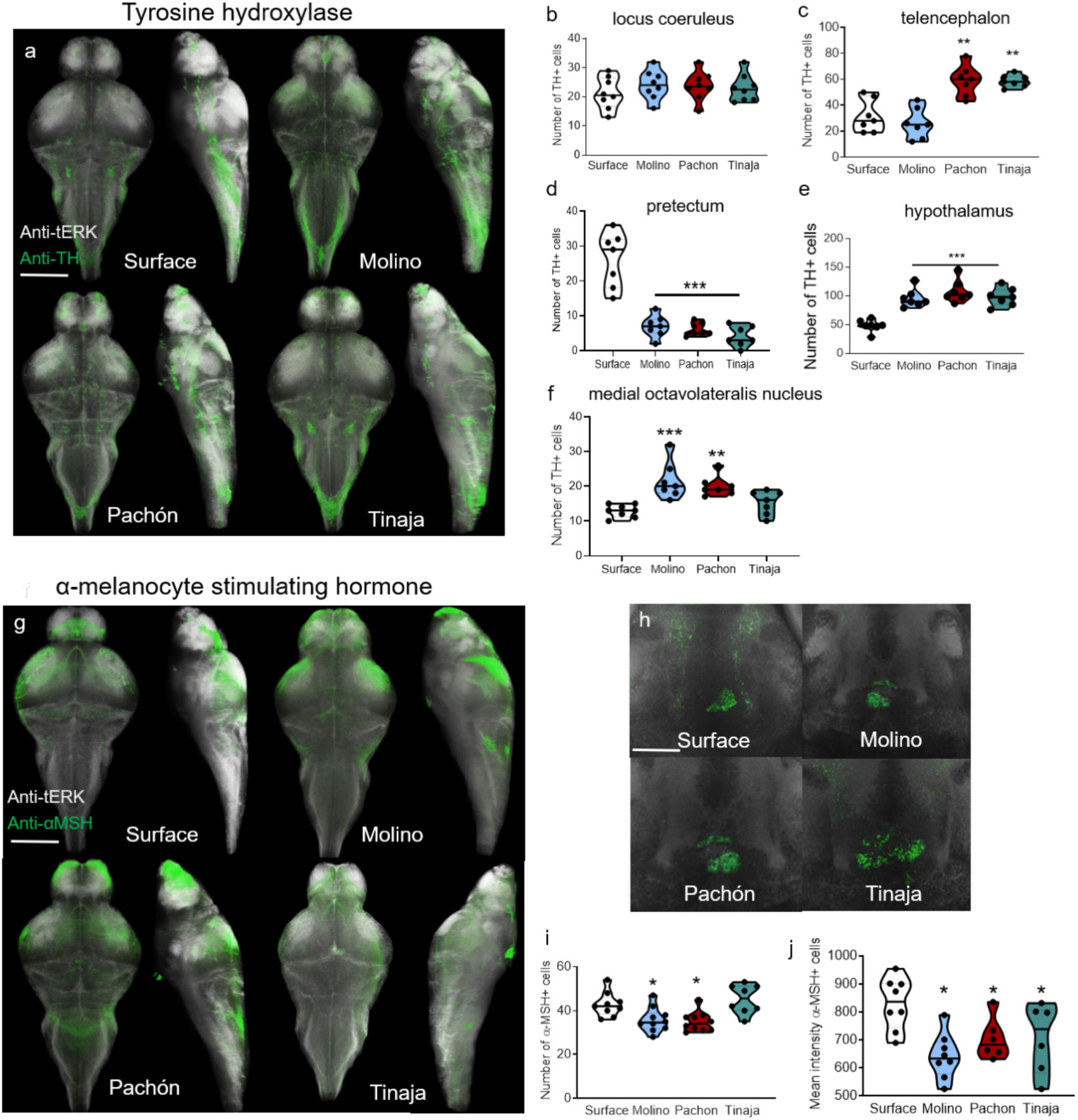
Whole-brain imaging of circuits associated with sleep and feeding. **A.** Whole-brain volumetric reconstructions of confocal imaging with anti-tERK (white) and anti-TH (green) for surface, Molino, Pachón, and Tinaja cavefish, with dorsal (Left) and sagittal (Right) views. Scale bar, 250 µm. **B-F** Numbers of cells expressing TH in distinct regions of the brain **B.** TH+ cell quantification in the locus coeruleus (1-way ANOVA, F=0.509, P>0.68, Molino, p>0.64, Pachon, p>0.54, Tinaja, p>0.83). **C.** TH+ cell number in the telencephalon (1-way ANOVA, F=18.87, P<0.001; Molino, p>0.66, Pachon, p<0.001, Tinaja, p<0.001). **D.** Quantification of pretectal TH+ neuron cells (1-way ANOVA, F=35.19, P<0.001; Molino, p<0.001, Pachon p<0.001, Tinaja, p<0.001). **E.** Number of TH+ cells in hypothalamus (1-way ANOVA, F=20.16, P<0.001; Molino, p<0.001, Pachon, p<0.001, Tinaja, p<0.001) **f.** TH+ cell number in the in the medial octavolateralis nucleus (1-way ANOVA, F=9.532, P<0.001; Molino, p<0.001, Pachon, p<0.002, Tinaja, p> 0.44) **G.** Whole-brain volumetric reconstructions of confocal imaging with anti-tERK (white) and anti-α-MSH (green) for four populations of *A. mexicanus*. Scale bar, 300 µm. **H.** Single-plane view of the α-MSH (green) cell cluster in the pituitary complex in surface fish and Molino, Pachón, and Tinaja cavefish. Scale bar denotes 50 µm. **I.** Total number of cells expressing α-MSH in pituitary complex **J.** Mean fluorescence intensity α-MSH+ individual cells from **i.** All comparisons were carried with n>8 and all posthoc tests compared cavefish to surface fish. N>8 for all measurements.

Finally, in the medial octavolateralis nucleus, a primary integration site of lateral line afferents, TH+ cell number was significantly greater in Molino and Pachón than in surface fish, but did not differ significantly between Tinaja and surface fish (Fig 2F, Fig S5). Together, these findings reveal that evolved differences in the number of neurons expressing TH can be independently regulated among distinct brain regions. In zebrafish, *hcrt*-expressing neurons localize to the rostral zone and pre-optic area of the hypothalamus and these neurons localized to similar regions across all *A. mexicanus* populations [50,52,53].

In all three cave populations, HCRT neurons were more abundant in both of these brain regions (Fig S6A-C), and the HCRT signal per cell was significantly elevated compared to surface fish (Fig S6D). A descending pathway along the midline that connects the midbrain to the spine stained strongly for HCRT in all cavefish populations, but not in surface fish (Fig S6E). In surface fish, and all three cavefish populations, HCRT-immunoreactive fibers localized to the locus coeruleus, as well as the lateral and intermediate zone of the hypothalamus, with ascending projections into the telencephalon (Fig S6E).

To determine whether cavefish evolved differences in neuropeptides that regulate feeding behavior, we examined the neuroanatomy of several conserved neuropeptides that regulate appetite. Genetic variants in the melanocortin receptor MC4R have been implicated in the regulation of feeding in diverse species, including *A. mexicanus* [19]. The neuropeptide α-melanocyte–stimulating hormone (α-MSH) antagonizes MC4R to inhibit feeding [54], and we identified an antibody that selectively label α-MSH neurons based on its known expression pattern. Immunostaining for α-MSH in surface fish is similar to that in zebrafish, predominantly labeling neurons in the pituitary complex with projections that ramify throughout the hypothalamus (Fig 2G) [55]. The number of α-MSH+ neurons was significantly reduced in both Molino and Pachón cavefish but did not differ between Tinaja and surface fish (Fig 2H). We identified differences in signal from α-MSH projections in a number of brain regions, including higher immunoreactivity in the cerebellum of surface fish than in all cavefish populations (Fig S7). In addition, the intensity of α-MSH signal from ascending projections that run laterally along the rostral zone of the hypothalamus into the forebrain was reduced in Pachón and Molino cavefish relative to surface fish (Fig S7), and the intensity of labeling within the tectum was reduced in all three cavefish populations. Together, the number of α-MSH neurons are reduced in the multiple cavefish populations, but not Tinaja, revealing differences in the evolution of feeding circuits.

The neuropeptide agouti-related protein (AgRP) opposes α-MSH signaling and functions as an inverse agonist of MC4R [54]. We found that AGRP localizes to the hypothalamus in surface fish and in all three populations of cavefish (Fig S8A), consistent with its expression in zebrafish [56], AgRP+ cells were more abundant in all three populations of cavefish than in surface fish (Fig S8B); in addition, the fluorescence intensity per cell was significantly higher in all populations of cavefish compared to surface (Fig S8C), indicating an increased number of AGRP+ cells and an increase in neuropeptide synthesis in cave populations. The projections of AgRP neurons shared many similarities with those of α-MSH neurons: AgRP+ fibers ran laterally in the medial hypothalamus cell bundle, with ascending fibers in the telencephalon forebrain bundle apparent in all populations (Fig S8D). We identified several significant differences in projections between populations, including tracts that connected the diffuse nucleus of the hypothalamus to the hindbrain in all three cavefish populations but were absent in surface fish (Fig S8D). Taken together, these findings are consistent with the adaptation of cavefish to a limited food environment and suggest changes in feeding circuitry that may underlie differences in prey-seeking behavior.

### Brain atlas reveals altered landscape of neural activity

The robust differences in behavior and neuroanatomy raise the possibility that brain activity differs between *A. mexicanus* populations. In zebrafish, phosphorylated ERK (pERK) accurately reflects neuronal activity with temporal resolution on the order of minutes [6]. To establish baseline differences in neural activity among *A. mexicanus* populations, we performed whole-mount immunostaining for pERK and tERK. *A. mexicanus* do not require food in their first week of life, and we collected non-fed fish between zeitgeber time (ZT) 4-6 (Where ZT 0 is the start of lights on (Fig 3A)). To assist in localizing pERK signal to distinct brain regions across multiple animals, we generated a common reference brain for each population using image registration (Fig S9A). To confirm accuracy of image registrations we implemented Jaccard image similarity analysis, which measures the volume of intersection between a registered brain and the template. This technique yielded a Jaccard index of 0.67 for surface, 0.71 for Molino, 0.69 for Pachón, and 0.70 for Tinaja, indicating that the registration algorithm was of high quality and that there was no significant variability among the populations (Fig S9B S Movie 3). Image registration revealed robust alignment of TH+ neurons in the locus coeruleus across populations of *A. mexicanus*, providing an average 3-dimensional positioning error of 5.6 µm in surface, 6.4 µm in Molino, 5.8 µm in Pachón, and 6.3 µm in Tinaja, comparable to published values in the zebrafish brain atlas [6] (Fig S9C,E). We then expanded the brain atlas to include all imaging data described above, including the sleep/wake regulating neurons expressing *hcrt* and *tyrosine hydroxlase* and the regulators of food consumption α-*msh* and *agrp*. We were able to generate a standard brain for four *A. mexicanus* populations, enabling us to directly examine markers of neural activity and defined neural populations within a single brain (Fig S9, S Movie 4).

**Figure 3:**
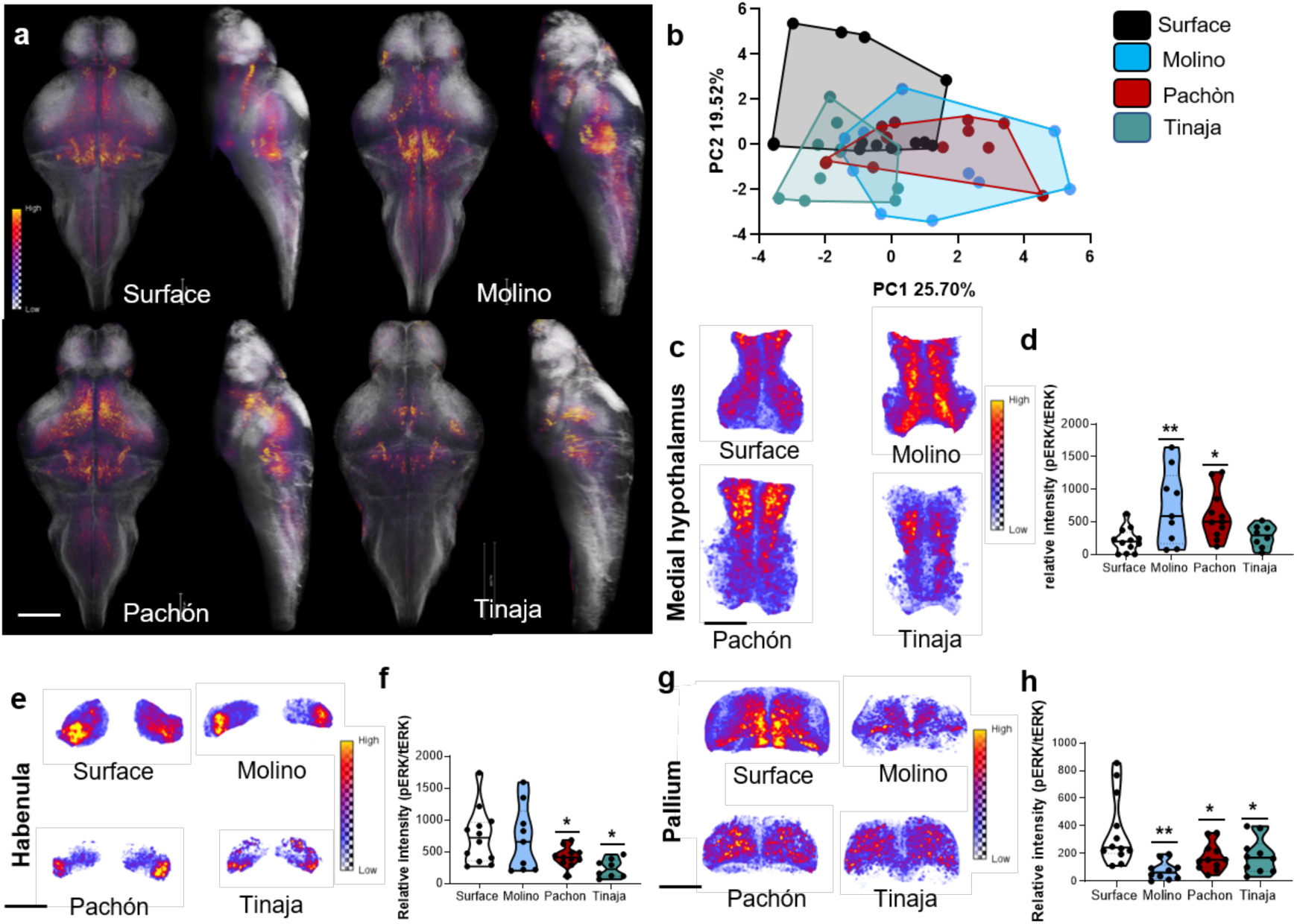
Whole-brain pERK neural activity imaging reveals altered landscape of brain activity. **A.** Average pERK activity maps overlaid onto standard brains with segmentations in major brain subdivisions in the indicated population of *A. mexicanus.* Scale bar denotes 200 µm. **B.** Principal component analysis of whole-brain neural activity in the brain of free-swimming fish. PC1 (one-way ANOVA, F=8.019, P<0.001; Dunnett post hoc, Molino, p<0.003, Pachón, p<0.001, Tinaja, p>0.95). PC2 (one-way ANOVA, F=8.786, P=0.0001; Dunnett post hoc, Molino, p<0.001; Pachón, p<0.05, Tinaja, p=0.001). Percentages indicate the amount of variance in neural activity explained by each PC. **C.** Maximum-intensity projections of mean pERK signal in medial hypothalamus. Scale bar denotes 100 µm **D.** Quantification of pERK signal in rostral zone of the hypothalamus. (one-way ANOVA, F=4.69, P<0.01; Molino, p<0.01, Pachón, p<0.04, Tinaja, p>=0.95). **E.** Maximum-intensity projections of pERK signal in habenula. Scale bar denotes 100 um. **f.** Quantification of pERK activity in the habenula (1-way ANVOA, F=4.16, P=0.012; Molino, p=0.99, Pachón, p=0.018, Tinaja, p<0.02). **G.** Maximum-intensity projections of pERK signal in the pallium. Scale bar denotes 100 µm **H.** Quantification of pallial neural activity (1-way ANOVA, F=6.18, P=0.001; Molino, p<0.001, Pachón, p<0.03, Tinaja, p<0.04). N>10 for all pERK activity mapping.

Next, we performed principal component analysis (PCA) on pERK/tERK imaging data to determine if distinct whole brain neural activity profiles have evolved in populations *of A. mexicanus*. The fish used in this experiment were actively moving prior to sample collection, and therefore considered to be awake. The PCA revealed distinctive clustering patterns of neural activity in each population. For example, Molino cavefish formed a cluster that was distinct from surface fish in both the first and second principal components (PC1 and PC2), whereas Pachón formed unique a cluster shifted to the right from surface fish in PC1 (Fig 3B). Tinaja cavefish formed a cluster below surface fish in PC2. PCA variable analysis revealed the candidate regions most strongly associated with altered neural activity in each principal component, including the rostral zone of the hypothalamus in PC1 (Fig S10A,B) and pallium and habenula in PC2 (Fig S10C). Collectively, PCA of pERK/tERK imaging from free-swimming *A. mexicanus* revealed distinct evolved patterns of neural activity within principal component space, suggesting cave-adapted fish have unique neural activity profiles compared to evolutionarily older surface fish.

To further characterize differences in neural activity among populations, we quantified the level of pERK activity in specific regions identified by PCA. As the rostral zone is thought to be homologous to the lateral hypothalamus in mammals, that serves as a critical regulator of both sleep and feeding behavior [57, 58], we speculated that activity within this region might differ between cavefish populations. Indeed, we observed a significant increase in pERK activity in rostral zone of the hypothalamus in Molino and Pachón cavefish populations relative to surface fish, but no differences between surface fish and Tinaja (Fig 3C-D). Furthermore, neural activity in the habenula, a region involved in stress response [59, 60], was significantly reduced relative to surface fish in Pachón and Tinaja, but unaltered? in Molino (Fig 3E-F). Finally, activity within the pallium, an area analogous to the mammalian amygdala and hippocampus that has been associated with emotion, motivation, and recently sleep regulation in zebrafish [61, 62], was significantly reduced in all populations of cavefish relative to surface fish (Fig 3G-H). We also quantified activity in 12 additional brain regions (Fig S11, Table 3). Together, this analysis reveals changes in brain regions associated with behaviors that have diverged between surface fish and cavefish.

### Neural activity associated with feeding behavior

To determine how brain activity differs during a multi-modal sensory behavior, we quantified the effects of feeding on neural activity. To this end, we compared brain-wide pERK levels in fish fed *Artemia* for 10 minutes that had not been fed and were freely moving prior to sacrifice (Fig 4A). We applied PCA to whole-brain activity patterns to determine whether feeding would create unique activity signatures in each population. Pachón and Tinaja cavefish formed distinct clusters in PC1 relative to surface fish, suggesting that they have evolved distinct neural activity patterns associated with feeding. By contrast, the evolutionarily younger Molino cavefish did not significantly differ from surface fish in either PC1 or PC2 (Fig 4B). PCA analysis revealed that brain regions clustered tightly in either PC1 (areas with differing responses among the populations) or PC2 (areas that exhibited variance among populations) (Fig S13A,B). The diffuse nucleus of the hypothalamus, was identified as the most significant variable in PC2, suggesting the hypothalamus integrates multimodal sensory inputs, which are activated by feeding in all populations (Fig S13C).

**Figure 4:**
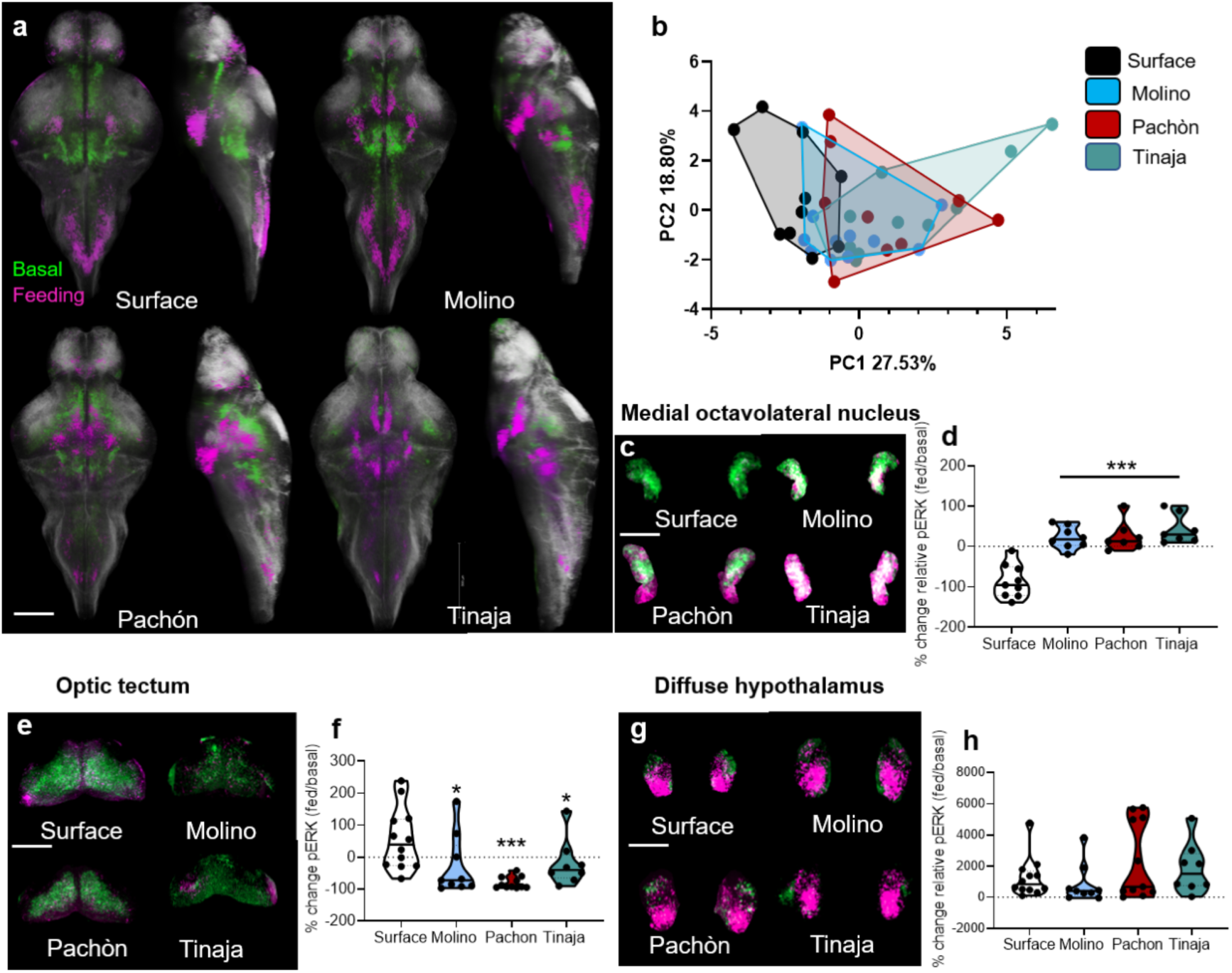
pERK neural activity during feeding reveals sensory transformation in cavefish and convergence on hypothalamic circuitry. **A.** Average whole-brain pERK activity patterns registered to standard brains (white) in non-feeding fish (green) and fish undergoing a 10-minute feeding assay (magenta). Scale bar is 200 *µ*m. **B.** PCA of whole-brain activity (reflected by pERK signal) in fish undergoing 10-minute feeding assay PC1 explained 27.53% of the variability of the brain activity (1-way ANOVA, F=6.652, P=0.001, Molino, p>0.40, Pachón, p<0.03, Tinaja, p<0.001). There were no statistical differences between populations? along PC2, which explained 18.80% of the variability. **C.** Maximum-intensity projection of pERK neural activity in medial octavolateralis nucleus activity (MON) for non-fed (green) and fed (magenta) fish. Scale bar denotes 50 *µ*m. **D.** Quantitation of the change in pERK activity in the MON during feeding (one-way ANOVA, F=22.14, P<0.001; Molino, Pachón, Tinaja p<0.001). **E.** Maximum-intensity projection of pERK activity in the optic tectum of non-fed (green) and feeding (magenta) fish Scale bar denotes 200 *µ*m **F.** Quantification of change in pERK activity in the optic tectum during feeding (one-way ANOVA, F=6.13, P=0.002; Molino, p>0.02, Pachon, p<0.001, Tinaja, p>0.04). **G.** Maximum-intensity projection of pERK activity in the diffuse nucleus of the hypothalamus of non-fed (green) and feeding (magenta) fish. Scale bar depicts 100 *µ*m. **H.** Quantification of change in pERK activity in the MON during feeding (1-way ANOVA, F=1.43, P=0.24; Molino, p>0.91, Pachon, p>0.33, Tinaja, p>0.8). N>10 for all feeding pERK neural activity.

All cavefish populations exhibited a significant increase in medial octavolateralis nucleus activity following feeding behavior, whereas surface fish exhibited a reduction, suggesting opposing polarities of medial octavolateralis nucleus activity during feeding between surface fish and all three cavefish populations (Fig 4C-D). Surface fish, like zebrafish, use visual cues to orient relative to prey. pERK level (i.e., neural activity) in the optic tectum was significantly higher in surface fish than in all three cavefish, suggesting that the tectum is not an input for feeding-associated behavior in cave populations of *A. mexicanus* (Fig 4E-F). Finally, feeding induced a robust increase in neural activity in the diffuse nucleus of the hypothalamus across all populations (Fig 4G-H), suggesting that this nucleus integrates multiple sensory modalities during feeding. Together, these findings highlight the evolution of brain-wide changes feeding behavior across multiple cave-adapted populations of *A. mexicanus*.

### Evolution of sleep-associated neural activity

Mapping brain activity during sleep in cavefish is difficult because individuals from these populations sleep for limited periods. However, the small size and relatively permeable blood brain barrier of *A. mexicanus* allows for measuring the effects of drugs on sleep regulation, similar to approaches for whole-brain imaging of sleep previously used in zebrafish [51,61,63]. Therefore, to directly compare brain activity in sleeping surface fish and cavefish, we pharmacologically induced sleep in all populations of cavefish. Previously, we showed that moderate concentrations of β-adrenergic antagonist propranolol and HCRT receptor inhibitor EMPA restore sleep to Pachón cavefish without affecting sleep in surface fish, suggesting enhanced sensitivity to inhibitors of β-adrenergic and HCRT signaling [44, 51]. The effects of these agents on neural activity and in additional cave populations is unknown.

Treatment with β-adrenergic antagonist propranolol and the HCRT receptor inhibitor EMPA restored sleep in all three cavefish populations, suggesting conserved signaling pathways contribute to sleep loss in independently-evolved cavefish populations (Fig 5A). Both drugs increased sleep bout length and bout number, without affecting waking activity, suggesting that the elevation of sleep is not due to lethargy (Fig S14). To determine whether the two drugs act induce similar or distinct changes in neural activity, we compared neural activity between awake DMSO-treated fish and sleeping fish treated with EMPA or propranolol (Fig 5B). Both EMPA and propranolol-treated sleeping surface fish and cavefish exhibited an overall reduction in neural activity compared to awake fish, similar to what was recently reported for drug-treated sleeping zebrafish (data not shown) [61]. In all populations, the results of PCA significantly differed between asleep/drug treated and and awake/DMSO treated fish, suggesting that pERK is a robust marker for detecting neural activity differences in a sleep-like state in *A. mexicanus* (Fig S15A).

**Figure 5:**
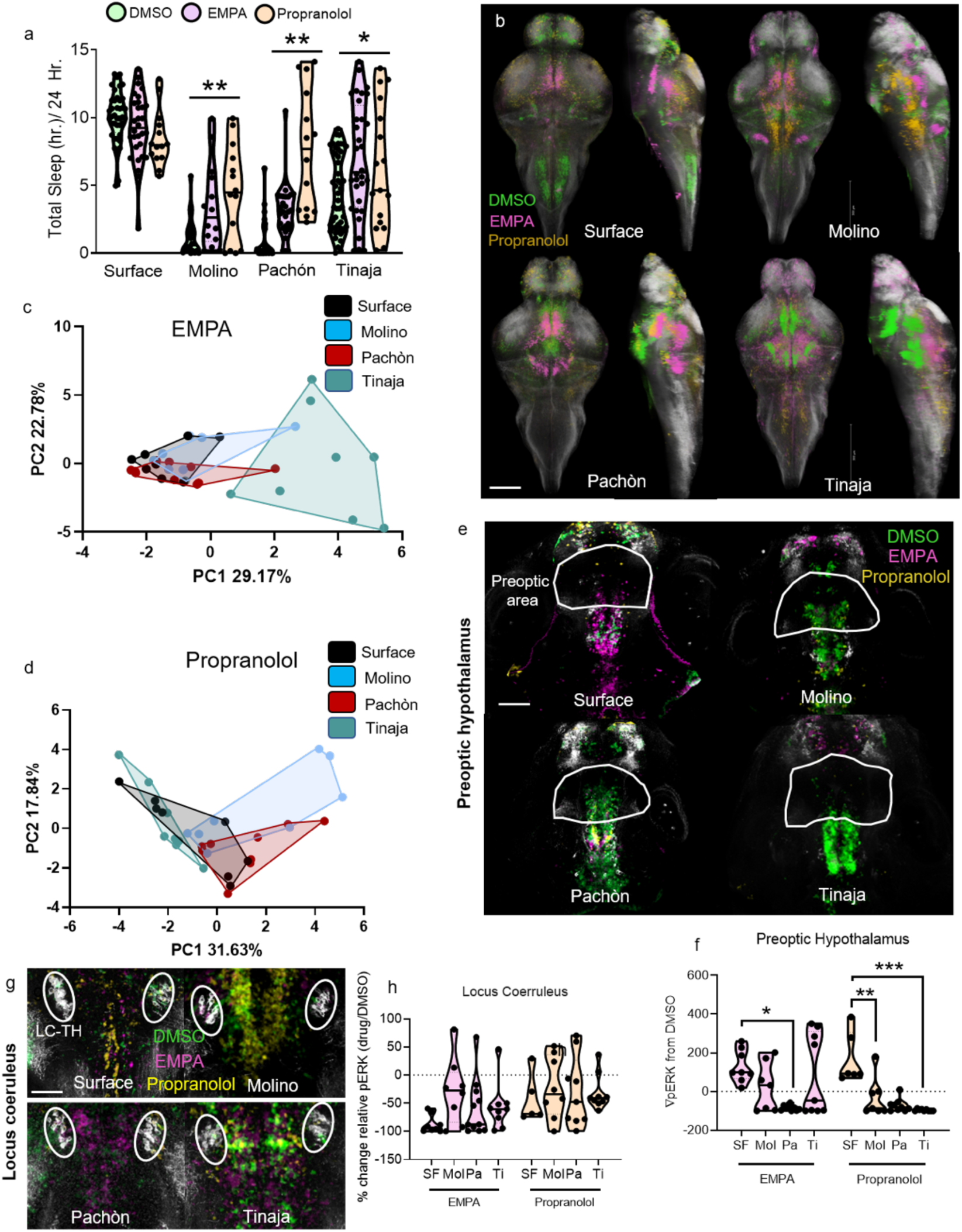
Whole-brain activity imaging of sleep-like state reveals heterogenous neural signatures. **A.** Quantification of 24 hour sleep recording with control DMSO (green) EMPA (magenta) and Propranolol (yellow) treatments (2-way ANOVA, F=34.50, P<0.001; Surface EMPA, P=0.894, Surface Propranolol, p<0.42, Molino EMPA, p<0.02, Molino Propranolol, p<0.01, Pachòn EMPA, p=0.011, Pachòn Propranolol, p<0.001, Tinaja EMPA, p<0.001, Tinaja Propranolol, p=0.034) **B.** Average whole-brain neural activity for surface, Molino, Pachón, and Tinaja cavefish under treatment with (waking) DMSO (green), (sleeping) EMPA (magenta), or (sleeping) propranolol (yellow). Scale bar denotes 200 µm. **C.** PCA of neural activity in fish treated with EMPA. PC1 explains 29.17% of the PCA variance (1-way ANOVA, F=16.75, P<0.001; Molino, p>0.56, Pachón, p>0.96, Tinaja, p<0.001). PC2 explains 22.78% of the variance (1-way ANOVA, F=0.849, P>0.47; Molino, p>0.84, Pachòn, p>0.74, Tinaja, p>0.90). D. PCA of neural activity in sleeping fish treated with propranolol PC1 explains 31.94% of the variation (1-way ANOVA, F=7.475, P<0.001; Molino, p<0.02, Pachón, p<0.05, Tinaja, p>0.61). PC2 explains 18.60% of the neural activity variation (1-way ANOVA, F=2.03, P=0.131; Molino, p=0.435, Pachòn, p=0.579, Tinaja, p=0.921). **E.** Single confocal plane view of average neural activity in preoptic area of the hypothalamus in awake DMSO (green) and sleeping EMPA (magenta) and sleeping propranolol (yellow). Scale bar denotes 100 µm. F. Quantification of the change in pERK neural activity in preoptic area of the hypothalamus in sleeping vs. waking fish (2-way ANOVA, F=6.959, P<0.001. For EMPA treatment: Molino, p>0.34, Pachón, p<0.001, Tinaja,p> 0.95; For propranolol treatment: Molino, p<0.02, Pachón, p<0.001, Tinaja, p<0.001). **G.** 20µm projection of hindbrain area containing TH+ locus coeruleus neurons (white circles) with average neural activity of awake DMSO (green) and sleeping EMPA-(magenta) or -propranolol-treated fish (yellow). Scale bar denotes 50 µm **H.** Quantification of the change in pERK signal (2-way ANOVA, F=1.71, P=0.124. EMPA: Molino, p=0.1, Pachòn, p=0.743, Tinaja, p=0.727; For propranolol treatment: Molino, p>0.97, Pachón, p>0.99, Tinaja,p> 0.99). N>10 for all pERK neural activity quantification, N>18 for all sleep behavior experiments.

Next, we sought to determine whether drug-treated fish in a sleep-like state converged upon shared or independent patterns of neural activity in each population. PCA analysis of EMPA-treated sleeping fish revealed that surface and Molino cavefish clustered tightly together, whereas Pachón cavefish formed a separate cluster in PC2 and Tinaja formed a separate cluster to the right in PC1 (Fig 5C). Variable analysis derived from PCA revealed that the main regions driving the changes along PC1 in Tinaja were in the telencephalon, including the pallium and subpallium, while the most significant variables for Pachón in PC2 were in the diencephalon, including several known sleep centers of the brain, such as the rostral zone and preoptic area of the hypothalamus (Fig S15B-D). Propranolol treatment also resulted in unique neural activity profiles across populations of *A. mexicanus*, with Molino and Pachón forming clusters of sleep-associated activity distinct both surface fish and Tinaja cavefish in PC1 (Fig 5D). PCA variable analysis revealed several highly-associated brain regions for both Propranolol and EMPA sleep conditions. These included regions that have been implicated in zebrafish or mammalian sleep regulation include the rostral zone and preoptic areas of the hypothalamus, and the locus coeruleus, indicating that shared regions of neural activity may be associated with sleep in *A. mexicanus* (Fig S15B-G).

We next quantified the changes in pERK activity in sleeping/drug-treated fish relative to waking DMSO-treated fish in different brain regions. In mammals, preoptic regions of the hypothalamus promote sleep [64]. Activity in the preoptic hypothalamus was robustly elevated in sleeping surface fish but was reduced or unchanged in all sleeping cavefish populations, revealing the presence of differentially evolved sleep signatures between surface and cave forms (Fig 5 E-F). In mammals and zebrafish, the locus coeruleus promotes wakefulness and receives inputs from wake-promoting HCRT neurons [47, 65]. We observed a significant reduction in neural activity in locus coeruleus TH+ neurons in all cave populations treated with either drug (Fig 5G-H). In sleeping fish, pERK activity was robustly elevated in a large area of the tegmentum, a sleep-promoting area in both mammals and zebrafish [59, 66]. In all populations, treatment with EMPA and propranolol increased tegmentum activity during sleep relative to DMSO-treated fish (Fig S16). In addition, activity was reduced during sleep across surface and all three cavefish populations in the rostral zone of the hypothalamus, a region containing HCRT neurons (Fig S16). Activity during sleep in numerous other regions, including the pallium, subpallium, intermediate zone of the hypothalamus, and cerebellum, differed among cavefish populations (Fig S16). Together, these results demonstrate unique activity signatures associated with sleep-like states across different *A. mexicanus* populations.

## Discussion

In this study, we used whole-brain imaging in fixed samples of independently evolved populations of *A. mexicanus*, an evolutionary model in which the cave and surface forms exhibit significant differences in complex behaviors, including sleep and feeding. Our systematic approach has revealed evolved alterations in neuronal organization at several levels, including morphology, circuitry, and neural activity. This work provides a basis for investigating the mechanisms by which evolution has altered brain morphology, and how these morphological changes are related to changes in behavior. In addition, our atlas will facilitate an unbiased examination of the relationship between the function and anatomy of different brain regions, as well as their relationship to the ecologies of each of the four populations studied.

Feeding behavior induced broad changes in brain activity across all four *A. mexicanus* populations, and likely activates brain regions associated with sensory processing, satiety, and motivation. While we identified differences in strike angle in two of the three populations of cavefish studied, it is possible that differences in brain activity or feeding-associated neurons are related to other aspects of feeding such as overall consumption, or vibration attraction behavior that emerge later in development. The evolution of sensory systems is particularly prominent in cavefish, including intrapopulation differences vision, mechanosensation, taste, and smell, in the regulation of behavior [67–70]. The identification of differences in anatomy and activity of numerous brain regions associated with the processing of sensory information, including the optic tectum and the medial octavolateralis nucleus, which receives information from the lateral line. Both of these regions were differentially active between surface and cavefish during feeding, suggesting that the two forms rely in different sensory modalities, or that these modalities are differentially processed.

We examined the effects of sleep-promoting drugs on brain activity in multiple *A. mexicanus* populations.. To date, sleep in *A. mexicanus* larvae and adults has been defined largely based on canonical methodology from zebrafish, which uses behavioral criteria such as quiescence and arousal threshold to define sleep [39, 40]. Recently, neural correlates of non-REM and REM sleep have been found in zebrafish using fluorescence-based polysomnography [61]. These studies localized synchronous activity associated with sleep to the dorsal pallium, raising the possibility that this region is analogous to the mammalian cortex [61]. Although the pERK imaging method used in this study does not have the temporal resolution to detect such events, we identified differences in neural activity within the dorsal pallium between *A. mexicanus* populations. The implementation of genetically encoded Ca^2+^ sensors in *A. mexicanus* will allow for greater temporal resolution of the differences in neural activity identified in this study, providing an opportunity to define sleep based on neural synchrony, similarly to methods commonly used in mammals.

The generation of whole-brain morphometric brain atlases enables localization of neuroanatomical regions associated with different behaviors [6,8,33]. To date, whole-brain atlases have been generated in fruit flies, zebrafish, and mice, allowing brains from different individuals or whole-brain Ca^2+^ imaging to be mapped onto a single standard brain [6,8,33,35,36,71–73]. The generation of these brain atlases in *A. mexicanus* represent the first use of whole-brain morphometrics to compare brain anatomy between different populations. This approach could be applied in model systems, including zebrafish and fruit flies, to identify differences in neuroanatomy between independent strains. For example, many behaviors differ between laboratory strains of zebrafish including stress, schooling, and feeding, [74–76], and in *Drosophila*, sleep and feeding behaviors differ among inbred populations [77, 78]. The generation of brain atlases for individual populations may provide insights into the neural mechanisms underlying these behavioral differences.

The development of a functional brain atlas in *A. mexicanus* will facilitate future efforts to better understand how evolution of the brain has led to behavioral divergence. Using individuals of the same species with a span of behavioral alterations will allow for direct interrogation of genotype–phenotype interactions; previously, such interactions have been difficult to parse by comparative approaches due to the relatively large phylogenetic divergences in other systems and the lack of functional tools. Isolated *A. mexicanus* populations represent diverse members of the same species, which is genetically amenable to transgenesis and mutagenesis techniques including the Tol2 transposase system and CRISPR/Cas9 engineering [38,79,80]. The brain atlas could be used as an anatomical marker to align whole-brain GCaMP imaging at a cellular resolution. The development of a functional brain atlas in *A. mexicanus* will facilitate future efforts to better understand how evolution of the brain has led to behavioral divergence. Isolated *A. mexicanus* populations represent diverse members of a single species, which is genetically amenable to transgenesis and mutagenesis techniques including the Tol2 transposase system and CRISPR/Cas9 engineering [38,79,80]. Here, our analyses in *A. mexicanus* is reliant on manual segmentation of brain regions, and our quantification consisted of 18 brain regions. In zebrafish, automated analysis, aided in part by increased resolution afforded by transgenic lines has allowed for segmentation into hundreds different brain structures [6–8]. The application of this technology, in combination with the use of genetically expressed anatomical marker, such as pan-neuronally expressed GCaMP has potential to compare the evolution of over 200 brain regions between populations.

Taken together these studies identify large scale differences between surface fish and cavefish populations of *A. mexicanus*, as well as between different populations or cavefish. This represents the first whole-brain anatomical brain atlas comparing intraspecies differences in brain structure and function. This resources has potential to provide information about the fundamental principles guiding the relationship between the evolution of brain function and behavior, as well as the contributions of naturally occurring variation in brain function that underlies behavioral differences between individuals.

## Materials and Methods

### Fish care

Animal husbandry was carried out as previously described [38, 81] and all protocols were approved by the IACUC of Florida Atlantic University. Adult breeding fish were housed in the university core fish facilities at a water temperature of 21±1°C. Lights were maintained on a 14:10 light-dark cycle throughout all experiments. Daylight intensity was between 25-40 Lux for both rearing and behavioral experiments. After nighttime breeding, larval fish were raised in an incubator at 23°C until 6 dpf to ensure consistent development. Fish were not fed until 6 dpf, and unless noted, fish were not in the fed state when sacrificed for imaging.

### Sleep behavior

Sleep behavior was assayed as previously described in [18]. Briefly, 6 dpf fish were individually housed in 24 well tissue-culture plates (Cat. No 662-102, CellStar) and acclimated for 18–24 hours before the beginning of the experiment at ZT0. Fish were recorded at 15 frames per second (fps) using a USB webcam equipped with a zoom lens and an IR-pass filter. Videos were saved as .avi files using the VirtualDub software, and then processed using the EthoVision XT (v12) behavioral profiling software. Raw locomotor data was exported as Unicode text, and then processed by custom-written code to calculate sleep parameters.

### Quantification of prey capture

At 6 dpf, larval fish were individually placed into circular wells with a diameter of 16 mm and a depth of 3 mm. After an acclimation period of 2 minutes, approximately 30 brine shrimp (*Artemia salina*) of the first instar stage were added to the well, and prey capture behavior was recorded from above at 100 fps for a period of 2 minutes. Recordings were acquired using a USB 3.0 camera (Grasshopper3, FLIR Systems) fitted with a zoom lens (75-mm DG Series Fixed Focal Length Lens, Edmund Optics Worldwide) and recorded with FlyCapture2 software (v2.11.3.163, FLIR Systems). To quantify prey capture dynamics, the angle of prey capture (strike angle) was measured for all successful feeding events in the 2-minute recording interval using the native “Angle” tool in ImageJ (NIH, v.1.51). All measurements were made in the frame prior to initiation of movement towards the prey. Strike angle was defined as the angle between the line segment extending down the fish’s midline and terminating parallel with the pectoral fins, and the line segment extending from this point to the center of the prey. Measurements of each strike were averaged to calculate the mean strike angle for that individual, and any recording with fewer than three feeding events was excluded from analysis.

### Quantification of brain activity during feeding behavior

Fish at 6 dpf were individually placed in 24-well plates (Cat. No 662-102, CellStar). After the fish were left undisturbed for 1 hour, approximately 30 brine shrimp were added to each well, and the fish were allowed to feed for 10 minutes. Fish were then immediately fixed in a 4% PFA solution and immunostained. Prior to immunostaining, feeding was visually confirmed based on the presence of brine shrimp in the gut of the fish.

### Pharmacology

All drug treatments were approved by the Florida Atlantic University IACUC committee (Protocols A15-32 and A16-04). For behavioral recording experiments, all fish were placed into individual wells of a 24-well plate and allowed to acclimate overnight before the beginning of the experiment. At ZT0, fish were treated with either solvent control, 0.1% DMSO, or freshly prepared propranolol (Sigma-Aldrich) or EMPA (Tocris Biosciences). Both drugs were dissolved in 100% DMSO, and then diluted to final concentrations of 0.1% DMSO and 30 µM propranolol or 100 µM EMPA. Behavior was then recorded for 24 hours across light-dark phases. For imaging experiments, all fish were treated with solvent, 0.1% DMSO, or freshly prepared propranolol or EMPA dissolved in DMSO. Fish were monitored from ZT2–ZT4 for bouts of inactivity associated with sleep (>60 seconds). Fish were sacrificed for imaging if they displayed a bout of inactivity of > 120 seconds. Control DMSO fish were sacrificed for imaging at any time after undergoing a swim bout during the 2-hour assay. Data are presented as DMSO (waking) and propranolol or EMPA (sleep-like) to characterize whole-brain activity under these unique conditions.

### Immunohistochemistry

Briefly, 6 dpf fish were strained through a plastic mesh sieve and then dropped into ice-cold 4% paraformaldehyde to kill them quickly before pERK activity resulting from sacrifice could be detected. The fish were fixed overnight, rinsed (here and below, rinses were in 0.3% PBT, performed three times for 15 minutes each), treated with 150 mM Tris-HCl (pH 9.0) for 15 minutes at 70°C, rinsed, incubated for 30 minutes on ice in 0.05% trypsin-EDTA, rinsed, placed in 3% H_2_O_2_ with 1% KOH for 15 minutes at RT to bleach pigmentation, and rinsed a final time. Fish were then placed in 0.3% PBT containing 2% DMSO, 1% BSA, and primary antibody at the indicated dilution: mouse anti-tERK (1:500), rabbit anti-pERK (1:500), rabbit anti-HCRT (1:500), rabbit anti-TH (1:500), sheep anti-α-MSH (1:5000), or rabbit anti-Agrp (1:400). Secondary antibodies were as follows: Alexa Fluor 488–conjugated anti-sheep Igg H+L, Alexa Fluor 488–conjugated anti-rabbit Igg H+L, and Alexa Fluor 561–conjugated anti-mouse Igg2a. See Table 2 for a complete list of concentrations and product numbers. Special care was taken when fish were being imaged for pERK, which is a fast indicator of neuronal activity; larval fish were sacrificed as quickly and consistently as possible and then processed essentially as described in [6].

### Image acquisition and analysis

All images were procured on a Nikon A1 upright confocal microscope equipped with a motorized piezo x-y-z stage and controlled by the Nikon Elements software. Fish were mounted dorsal side up in 2% low–melting temperature agarose (Sigma A9414) on a microscope slide (Fisher 12-518-101) in a glass-bottomed chamber. Individual fish were held in 100–150 µL agarose. A tiling function was used to image the entire brain and imaged were stitched together both images with a 15% overlap on the x-y plane. All images were acquired at 2-µm steps. Cell counts and intensity were quantified using the Nikon Elements software (4.5). Individual regions of interest (ROIs) were drawn over each detected cell. Mean intensity was calculated by subtracting background intensity, extracting the entire stack signal into Excel, and then restricting the quantified signal to ROIs matching the cells. To segregate neuronal populations within anatomical regions, brains were registered and overlaid with the label field. Cells within specific nuclei were then quantified within that region.

### Morphometric analysis

All morphometric image analysis was performed using the FEI Amira software. Confocal stacks were imported into Fiji/ImageJ (1.52), then imported to Amira (6.2.1). A mask was applied to include only neural tissue in the field of view. Brain regions were then manually segmented using the “lasso tool” with automatic edge detection. A developmental map was created that included the main divisions of the brain, including spine, rhombencephalon, mesencephalon, diencephalon, and telencephalon. These large divisions were segmented with tERK antibody, which defines physical divisions between regions. This result was saved as a label field for both the template brains, as well as for each animal that was segmented. A second anatomical map of smaller nuclei was generated, including cerebellum, optic tectum, optic neuropil, habenula, pineal gland, rostral zone of the hypothalamus, diffuse nucleus of the hypothalamus, intermediate zone of the hypothalamus, pre-optic nucleus, pallium, sub pallium, and pituitary complex. For the template brain of each population, six different cell markers (anti-tERK, anti-TH, anti-HCRT, anti-αMSH, anti-AgRP, and anti-pERK) were used to guide segmentation of regions by expression pattern. The template maps for each population were then used to guide all other segmentations. The material statistics, containing all raw data regarding sizes and locations of regions, were then exported, and percent of brain volume was calculated for each region. Data was analyzed in MATLAB (2019b) or GraphPad (v8) and visualized by ranking regions by size and graphing in a MATLAB ribbon plot. Statistical differences in region size were determined by performing one-way ANOVA with a Dunnett post-hoc test to detect changes in region volume across populations.

### Image registration

A template brain for each population of *A. mexicanus* was imaged using tERK immunological stain to label all neural tissue. The voxel size for the template was 0.61×0.61 × 2 µm^3^ (x × y × z). The template and transformation brains were loaded into Amira, and the “Register images” module was loaded. The transformation model included 12 degrees of freedom to account for ridged, isotropic, anisotropic, and shearing transformations. The outside threshold was set to 0.8. A correlation metric was used as the model for the transformations. A histogram filter was applied between 100-4095 to remove dark background pixels from the transformation calculation, which significantly reduced registration time. The coarsest resampling rate was set to either 16 × 14 × 6 or 14 × 16 × 6 depending on the reference brain used, and “ignore finest resolution” was unchecked to increase registration accuracy. Population averages of each antibody stain were generated by loading registered stacks into Amira and selecting the “average volumes” module. A single image stack was then made to represent the average expression pattern of three to ten individuals for each antibody stain from each population.

### pERK activity mapping

All analyses were performed in FIJI/ImageJ, Amira, and Matlab. After registration, the pERK channel was divided by the tERK channel. Histogram restoration was performed to restore the original span of pixels acquired on the confocal. A gaussian filter was then applied to smooth pixels, with pixels saturation set to 70% of the maximum pixel intensity and cut at 5% of the peak lowest pixel intensity to delete background noise, similar to previous analysis in zebrafish [6]. This range of pixels generated a cell mask of pERK-positive cells and removed background from quantitative analysis (Fig 3S3). Individual image stacks were quantified by extracting all voxels and determining the mean signal of voxels per anatomical region denoted within the template brains.

### Principal component analysis

Using Principal Component Analysis (PCA), we can consider each individual as a coordinate in a space whose axes are linearly-independent combinations of regional brain activity ranked according to total inter-individual variance of their activity as characterized by pERK expression. We find that the first two principal components (PC) capture 47.3%-58% of all activity variance across brains, and given the exploratory nature of this work, we are satisfied to begin here, especially considering the otherwise less comprehensible task of quantifying these to higher order, which is beyond the scope of this work. We thus transform the eighteen-dimensional activity space into just two-dimensions where each PC (dimension) comprises a combination of brain regions grouped by their alignment (correlation) with each other and ranked according to the magnitude of their variance. Principal component analysis was performed in R and XLStat.

All pERK voxels were isolated by brain regions, including developmental regions, as well as all smaller regions, resulting in 20 different components for the PCA. The first two components accounted for between 47.3% - 58% of the total variability across all brains. Statistical differences between populations by PCA were detected using a 1-way ANOVA with posthoc analysis.

### Generation of brain atlas

The standard brain for each population was generated by registering all brains with tERK to a tERK+ template brain, with a separate template for each population. Each brain represents a unique transformation to align to the template. Thus, individual brains were registered, and then the transformation matrix generated by the tERK channel was applied to the second channel, which imaged the protein of interest. Briefly, the steps in Amira were as follows. Population averages for each protein marker were calculated from between 3 and 18 fish. To generate the average, the “average volumes” module was loaded, and then the transformed stacks were loaded, with the resultant single image representing the average of all images processed. This was performed for each population for HCRT, TH, AgRP, α-MSH, and pERK for baseline conditions and feeding or drug treatments. Each of these average stacks were saved to represent the average expression pattern for that protein for each population. If any expression outside the brain was present in a stack, it was excluded by generating a mask to delete it from view.

## Acknowledgements

We are grateful for feedback and helpuful discussion from many groups in the zebrafish community including Harry Burgess (NIH) German Sumbre (IBENS) and Robert Kozol (FAU). This work was supported by NIH awards R21NS105071 and 1R01GM127872 to ACK; NSF awards IOS 165674 to ACK, DEB 174231 to ACK and JEK, and an NSF EDGE award 1923372 to ACK and ERD; and US-Israel BSF award SP#2018-190 to ACK and LA.

## Supplemental Figures

**Supplementary figure 1: Quantification of sleep architecture in individual *A. mexicanus* populations. A.** Waking activity did not vary significantly among the four populations of *A. mexicanus* (1-way ANOVA, F=0.518, P=0.672; Molino, p>0.61, Pachón, p>0.99, Tinaja, p>0.81). **B.** All cave populations have converged upon significant increases in total locomotion per 24 hours relative to surface fish (1-way ANOVA, F=13.45, P<0.001; Molino, Pachón, Tinaja, p<0.001). **C.** Average sleep bout duration was significantly reduced in all populations of cavefish compared to surface fish (1-way ANOVA, F=35.82, P<0.001; Molino, Pachón, Tinaja, p<0.001). **D.** Sum of bout number was significantly reduced in all cave populations relative to surface fish (1-way ANOVA, F=22.01, p<0.001; Molino, Pachón, Tinaja, p<0.001).

**Supplementary figure 2: Whole brain size does not differ among populations of *A. mexicanus.* A.** Whole-brain confocal scans of tERK antibody signal. Each population is depicted in dorsal (left) and sagittal (right) aspects. Scale bar denotes 300 µm. **B.** Quantification of whole-brain scans revealed no significant difference among populations in brain size (1-way ANOVA, F=0.869, P=0.46; Molino, p>0.69, Pachón, p>0.27, Tinaja, p>0.59).

**Supplementary figure 3: Evolution of neural developmental regions. A.** Volumetric projection of diencephalon (magenta). **B.** Convergence of expanded diencephalon in cavefish populations (1-way ANOVA, F=3.56, P=0.02; Molino, p<0.02, Pachón, p<0.01, Tinaja, p<0.04). **C.** Volumetric projection of mesencephalon (cyan). **D.** Reduction of mesencephalon in all cavefish populations (1-way ANOVA, F=26.72, p<0.001; Molino, Pachón, Tinaja, p<0.001). **E.** Volumetric projection of rhombencephalon (red). **F.** Rhombencephalon is expanded in cavefish relative to surface fish (1-way ANOVA, F=15.15, p<0.001; Molino, p<0.01, Pachón, p<0.001, Tinaja, p<0.001). **G.** Volumetric projection of telencephalon (green). **H.** Telencephalon size did not significantly differ among *A. mexicanus* populations (1-way ANOVA, F=0.845, P=0.47; Molino, p>0.42 Pachón, p>0.35 Tinaja, p>0.68). Scale bar for all images is 300 µm.

**Supplementary figure 4: Evaluation of neuroanatomical morphology in *A. mexicanus.*** Change in size of each region (as a percentage of the whole brain) relative to surface fish. See Table 1 for details and statistics.

**Supplementary figure 5: Neuroanatomical characterization of the TH circuitry.** Close-up views of cell clusters within nuclei across the brain in surface, Molino, Pachón, and Tinaja. Left, locus coeruleus; second from left, telencephalon; middle, hypothalamus; second from right, pretectum; right, medial octavolateralis nucleus.

**Supplementary figure 6: Convergent evolution of HCRT in cave-dwelling *A. mexicanus.* A.** Whole-brain volumetric projections of tERK (white) and hypocretin (green). Scale bar, 300 µm. **B.** Quantification of hypocretin cells in rostral zone of the hypothalamus in 6-day old fish reveals a convergence of enhanced HCRT in all cave populations relative to surface fish (1-way ANOVA, F=22.5, P<0.01; Molino, Pachón, Tinaja, p<0.001). **C.** Preoptic hypothalamus HCRT cluster was significantly larger in cave-adapted populations than in surface fish (1-way ANOVA, F=9.21, P<0.001; Molino, Pachón, Tinaja, p<0.001). **D.** Fluorescence intensity per HCRT cell was higher in cavefish in both rostral zone of the hypothalamus preoptic hypothalamus (1-way ANOVA, F=20.69, P<0.001; Molino, Pachon, Tinaja, p<0.001).

**Supplementary figure 7: Reduced number of α-MSH neurons in cavefish.** Single-plane views of confocal scans showing tERK (white) and α-MSH staining across the brain of 6-dpf *A. mexicanus.* Left panel: the cerebellum exhibits greater immunoreactivity in surface fish than in all populations of cavefish. Second from left: hindbrain expression was largely the same across all populations of *A. mexicanus.* Second from right: optic neuropil was highly immunoreactive in surface fish, with lower expression in all cave populations. Right: telencephalon was highly immunoreactive to MSH in surface and Tinaja, but not in Molino or Pachón.

**Supplementary figure 8: Increased number of AgRP neurons in cavefish. A.** Whole-brain volumetric projections of tERK (white) and AgRP (green). Scale bar, 300 µm. **B.** Quantification of AgRP+ cells in the pituitary complex reveals a convergence on higher numbers of cells in all populations of cavefish relative to surface fish (1-way ANOVA F=11.18, P<0.001; Molino, p<0.02, Pachón, p<0.02, Tinaja, p<0.001). **C.** Fluorescence intensity did not differ among populations of cavefish or surface fish (1-way ANOVA, F=0.88, P=0.46; Molino, p>0.82, Pachón, p>0.70, Tinaja, p>0.97). **D.** Single-plane confocal views of AgRP expression in the ventral hypothalamus, showing immunoreactive fibers in the hypothalamic and forebrain bundles (left). In the medial plane of the hypothalamus, signal was intense in the diffuse nucleus of the hypothalamus in Molino and Pachón, but not in surface or Tinaja (middle). All cave populations exhibited intense IR expression at the midline through the hindbrain. No such expression was observed in surface fish (right).

**Supplementary figure 9: Image registration in *A. mexicanus* brains. A.** Examples of brains before (top) and after (bottom) alignment to the template brain (green) and transformed brain (green) for four populations of *A. mexicanus.* **B.** Jaccard image similarity analysis detected no differences in registration quality across populations. (1-way ANOVA, F=0.02, P=0.99). **C.** Registrations applied to an anatomical label (anti-TH). Left panel shows TH staining in locus coeruleus for three fish, each colored differently. Right panel shows population average of TH+ expression in locus coeruleus from 10 fish. **D.** Registered labels were combined together to create the standard brain and applied to morphological neuroanatomy. Means of six different labels and five different segmented areas are shown. **E.** Mean distance error of registered TH+ cells in the locus coeruleus was quantified for all populations. Error did not differ significantly across populations (1-way ANOVA, F=2.27, P=0.09).

**Supplementary figure 10: Variable analysis for anatomical contributions to PC1 and PC2. A.** Variable vectors revealed a spread of regions affecting both PC1 along the x-axis and PC2 along the y-axis. **B.** Top five variables that contribute to PC1, including rhombencephalon, diencephalon, locus coeruleus, rostral zone of the hypothalamus, and preoptic region of the hypothalamus. These five regions contribute to 55.09% of the variability in PC1. **C.** Top regions for variation in PC2, including the telencephalon, optic tectum, habenula, pallium, and subpallium. Together, these regions contribute 66.31% of the variation in PC2.

**Supplementary figure 11: Quantification of neural activity in developmental divisions of the brain. A.** Max intensity projection of pERK activity in the telencephalon. **B.** Reduced forebrain activity in cave-adapted populations (1-way ANOVA, F=4.78, P=0.005’ Molino, p<0.003, Pachón, p=0.06, Tinaja, p<0.01) **C.** Max intensity projection of diencephalon pERK activity. **D.** Quantification of pERK in diencephalon (1-way ANOVA, F=7.51, P<0.001; Molino, p<0.03, Pachón, p<0.03, Tinaja, p=0.2). **E.** Max intensity projections of neural activity measured by pERK in the mesencephalon. **F.** Quantification of mesencephalon neural activity (1-way ANOVA, F=2.61, P=0.063; Molino, p>0.13, Pachón, p>0.99, Tinaja, p>0.86) **G.** Rhombencephalon pERK activity by max projection. **H.** Quantification of neural activity in hindbrain (1-way ANOVA, F=3.82, P>0.01; Molino, p>0.28, Pachón, p>0.08, Tinaja, p>0.69). Scale bar denotes 200 µm for all images.

**Supplementary figure 12: Pipeline for pERK quantification for whole-brain activity mapping. A.** Top left panel: single plane confocal scan with tERK (white) and pERK (green). Top right panel: pERK/tERK with brain mask applied after registration to the template brain. Bottom left: application of cell mask; blue represents cells that were kept for the activity map that were statistically significantly different from background. Bottom right: Activity map (green) applied to the template brain (white). **B.** pERK/tERK ratio histograms for all basal free-swimming fish. Cell mask was generated for each individual fish by applying a 5% cutoff from the peak of the histogram.

**Supplemental figure 13: PCA and variable analysis for feeding brain activity. A.** Loading plot. **B.** PC1. **C.** PC2

**Supplementary figure 14: Sleep architecture for Propranolol and EMPA treatments. A.** Waking activity is not altered by drug treatments (2-way ANOVA, F=1.43, P>0.2) **B.** Average bout duration was altered by EMPA and Propranolol treatment in *A. mexicanus* (2-way ANOVA, F=20.04, P<0.001; EMPA: Surface, p>0.82, Molino, p>0.02, Pachón, p>0.04, Tinaja, p>0.03. Propranolol: Surface, Molino, p>0.03, Pachón, p>0.001, Tinaja, p>0.12) **C.** Total sleep bout number was altered by both EMPA and Propranolol treatments (2-way ANOVA, F=48.77, P<0.001; EMPA, Surface, p>0.85 Molino, p>0.04 Pachón, p>0.02 Tinaja, p>0.07. Propranolol: Surface, p>0.94 Molino, p>0.02 Pachón, >0.001 Tinaja, p>0.01).

**Supplementary figure 15: PCA and Variable analysis for anatomical contributions to PC1 and PC2 during drug treatments. A.** Loading plot for EMPA treatment **B.** Top anatomical regions from variables in PC1 for EMPA treatment **C.** Top neuroanatomical regions for PC2 in EMPA treatment **D.** Loading plot for Propranolol treatment. **E.** Top neuroanatomical regions for PC1 in Propranolol treatment **F.** Top neuroanatomical regions for PC2 during Propranolol treatment.

**Supplementary figure 16: Regions with sleep-associated changes in neural activity. A.** Rostral zone of the hypothalamus (2-way ANOVA, F=2.24, P<0.04; EMPA: Molino, p>0.99, Pachón, p>0.61, Tinaja, p>0.94. Propranolol: Molino, p>0.99, Pachón, p>0.99, Tinaja, p>0.99) **B.** Tegmental area (2-way ANOVA, F=0.38, P>0.91; EMPA: Molino, p>0.93, Pachón, p>0.99, Tinaja, p>0.92. Propranolol: Molino, p>0.99, Pachón, p>0.99, Tinaja, p>0.98 **C.** Pallium (2-way ANOVA, F=2.24, P<0.04; EMPA: Molino, p>0.99, Pachón, p>0.61, Tinaja, p>0.94. Propranolol: Molino, p>0.99, Pachón, p>0.99, Tinaja, p>0.99). **D.** Subpallium (2-way ANOVA, F=10.01, P<0.001; EMPA: Molino, p>0.99, Pachón, p>0.99, Tinaja, p<0.001. Propranolol: Molino, p>0.99, Pachón, p>0.76, Tinaja, p>0.55) **E.** Intermediate hypothalamus (2-way ANOVA, F=4.00, P<0.001; EMPA: Molino, p<0.02, Pachón, p>0.001, Tinaja, p>0.01. Propranolol: Molino, p>0.95, Pachón, p>0.24, Tinaja, p>0.05). **F.** Optic tectum (2-way ANOVA, F=8.55, P<0.001; EMPA: Molino, p>0.99, Pachón, p>0.99, Tinaja, p>0.89. Propranolol: Molino, p>0.04, Pachón, p<0.001, Tinaja, p>0.99).

**Supplemental Video 1:** Representative neuroanatomical segmentations through whole brain.

**Supplemental Video 2:** Three-dimensional reconstructions of neuroanatomical segmentations for surface, Molino, Pachón, and Tinaja fish. tERK antibody was used to visualize neural tissue. Colors denote anatomical regions, which were labelled computationally.

**Supplemental Video 3:** Example stacks showing transformed brains (magenta) and template brains (green).

**Supplemental video 4:** Standard brain for each population of *A. mexicanus.* Neural tissue is visualized with tERK antibody (white). Anti-TH (green) anti-HCRT (magenta) anti-AgRP (red) anti-MSH (cyan). Average pERK activity during basal conditions in orange.

**Supplemental video 5: Basal whole brain activity throughout whole brain.** pERK activity is shown in green, registered onto template brains (white).

**Supplemental video 6: Whole-brain activity during non-fed and fed conditions.** pERK neural activity for non-fed (green) and fed (magenta) aligned to reference brains in tERK (white) for each population.

**Supplemental Video 7: Whole-brain activity during drug-induced sleep.** Neural activity associated aking DMSO (green) and sleeping EMPA (magenta) and Propranolol (yellow)

**Supplemental Table 1.** List of segmented regions with sizes and statistics.

**Supplemental Table 2:** Antibodies used with concentrations and part numbers

**Supplemental Table 3:** Basal pERK/tERK neural activity during basal conditions

**Supplemental Table 4:** Neural activity measured with pERK/tERK activity during feeding.

**Supplemental Table 5:** pERK quantification in all segmented brain regions during drug treatments.

